## Supplementary Figures for "Diverse and abundant phages exploit conjugative plasmids"

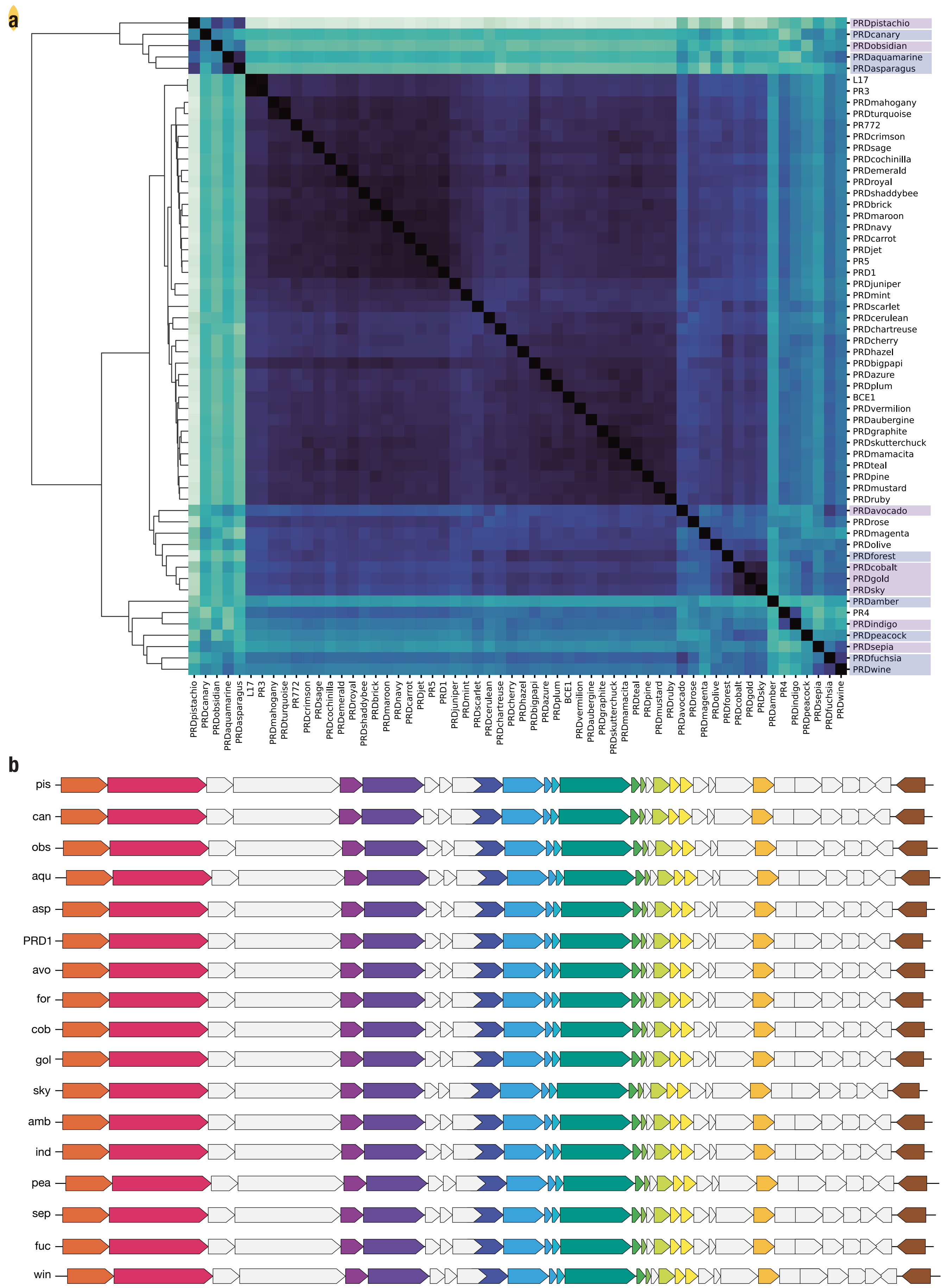


**Fig S1 | a,** Whole-genome pairwise nucleotide identity matrix comparing all known alphatectiviruses. Highlighted isolates represent proposed new species. **b,** Gene map comparison of representative alphatectiviruses isolated in this study, colors are as in (Figure S6b). Chosen isolates are those that meet species demarcation criteria according to current guidelines by the International Committee on Taxonomy of Viruses (ICTV) (<95% average nucleotide identity).

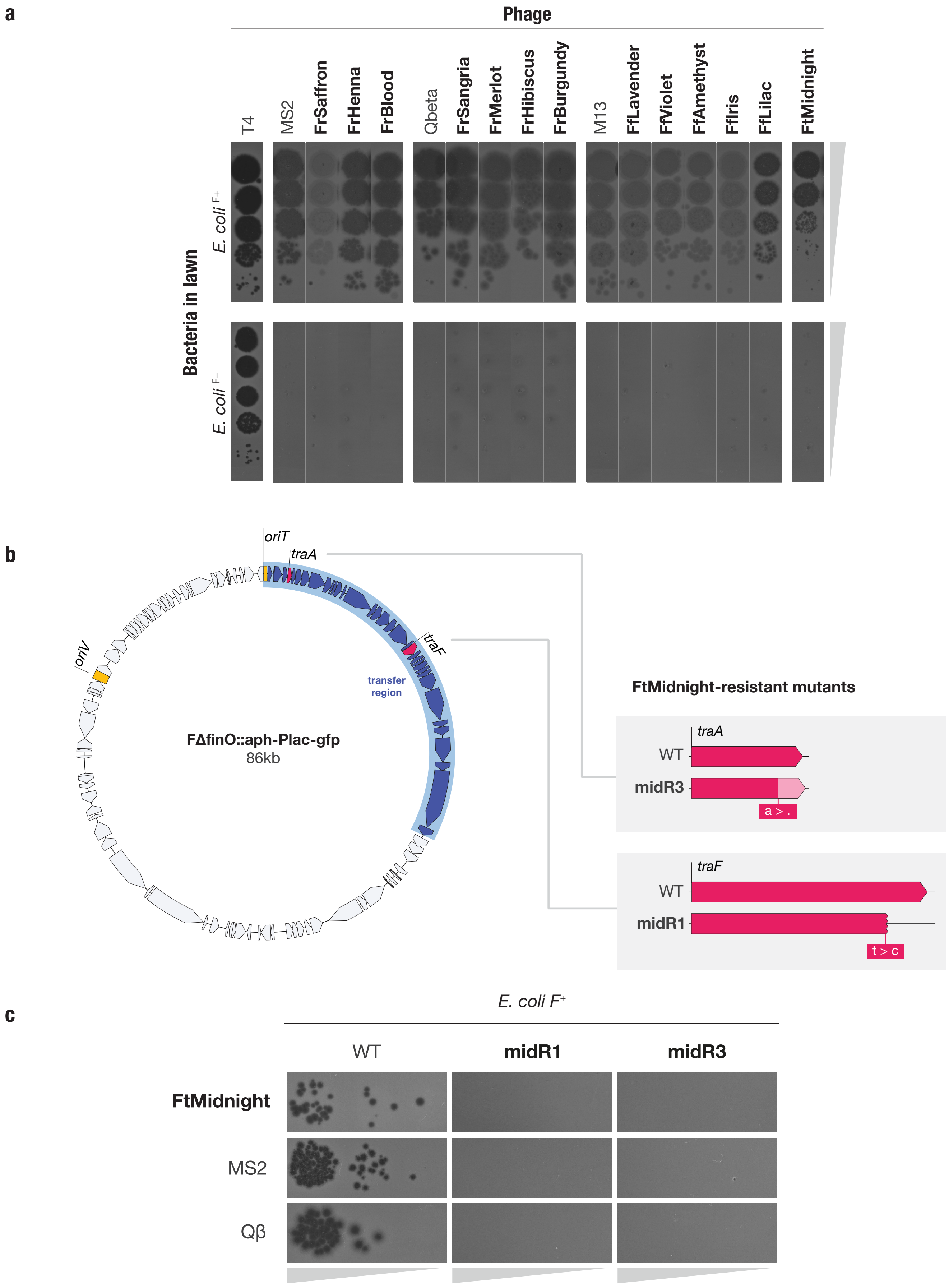


**Fig S2 | a,** Confirmatory plaque assay of all newly isolated F dependent phages on *E. coli* MG1655 hosts with and without the F plasmid, confirming plasmid dependency. The F plasmid derivative FΔfinO::aph-Plac-gfp is used throughout this work (Methods). *E. coli* phage T4, which is not plasmid-dependent, is included as a control **b,** Mutations conferring resistance to FtMidnight map to the transfer region of the F plasmid. Independently, mutant midR3 has a 1 bp deletion that causes a frameshift in the C-terminal domain of TraA, the major pilin, resulting in missense amino acid sequence at the C-terminus of the protein. Mutant midR1 has a substitution that causes a premature truncation of protein TraF, which is an outer-membrane component of the plasmid transfer machinery. Both proteins are required for pilus assembly and plasmid transfer, suggesting that FtMidnight interacts directly with the conjugative pilus. **c,** F-plasmid mutations that confer resitance to FtMidnight (midR1 and midR3) confer collateral resistance to F-plasmid dependent phages MS2 and Qbeta.

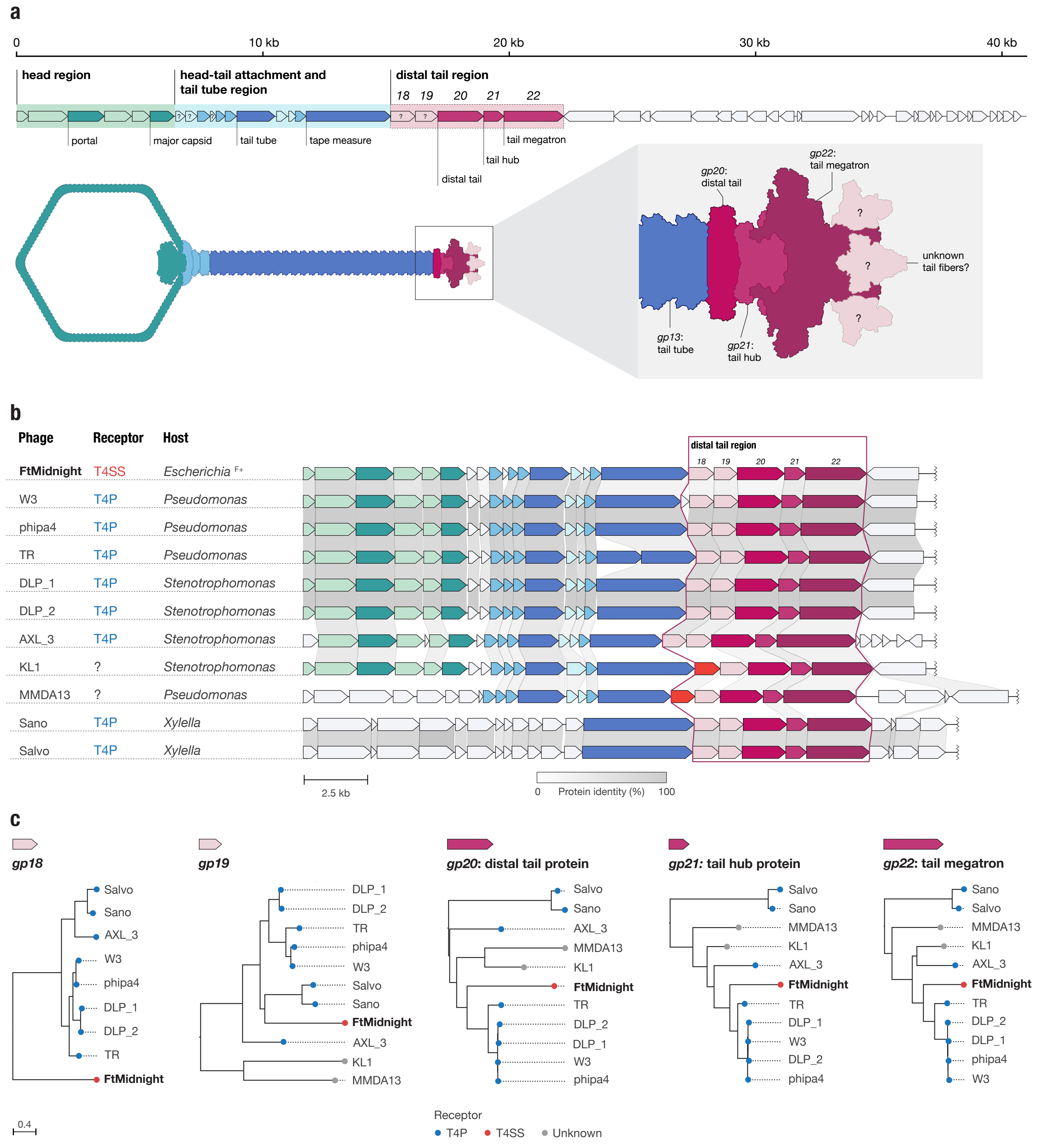


**Fig S3 | a,** Gene map and structural model of FtMidnight. The model was constructed through annotation with the PHROGs database, and refined with structure-guided homology matches to PDB structures using HHpred (Methods). The predicted head formation region is highlighted in green, the predicted head-tail attachment and tail tube region is highlighted in blue, and the genes predicted to be involved in the distal tail formation (gp18-22) are highlighted in pink. The distal tail region is presumed to interact with the receptor in the bacterial cell surface. Genes of unknown function within those regions are marked with a question mark. In the distal tail region, genes gp20-gp22 match components of the PDB structure 8GTC, which is the distal tail region of the marine siphophage vB_DshS-R4C. (See Table S2). Homologs to the tail fiber in vB_DshS-R4C were not identified. **b,** Comparison of phage genomes containing homologs of gp18-gp22. Homologous genes are color-coded and aligned, and the connector shade corresponds to protein identity. Distal tail region is framed to show conservation of the region. All phages have perfectly conserved gp18-22, except KL1 and MMDA13, which have a distinct gene, shown in orange, in place of gp18. Host of isolation and characterized phage receptor is shown on the left. While FtMidnight uses a type 4 secretion system (T4SS), as a receptor (the F-pilus structure, in particular), the rest of the characterized phages shown use the type 4 pilus (T4P). The receptor was annotated if found in the literature. Accessions and references can be found in Table S2. **c.** Phylogenetic trees of individual proteins of the distal tail region from the phage genomes shown in (b). Leaves in the trees are colored based on the known receptor for the phage of origin. gp18 appears to be the most divergent of the distal tail region proteins from homologs in T4P-associated phages, suggesting it might be responsible for the difference in receptor specificity.

**a**

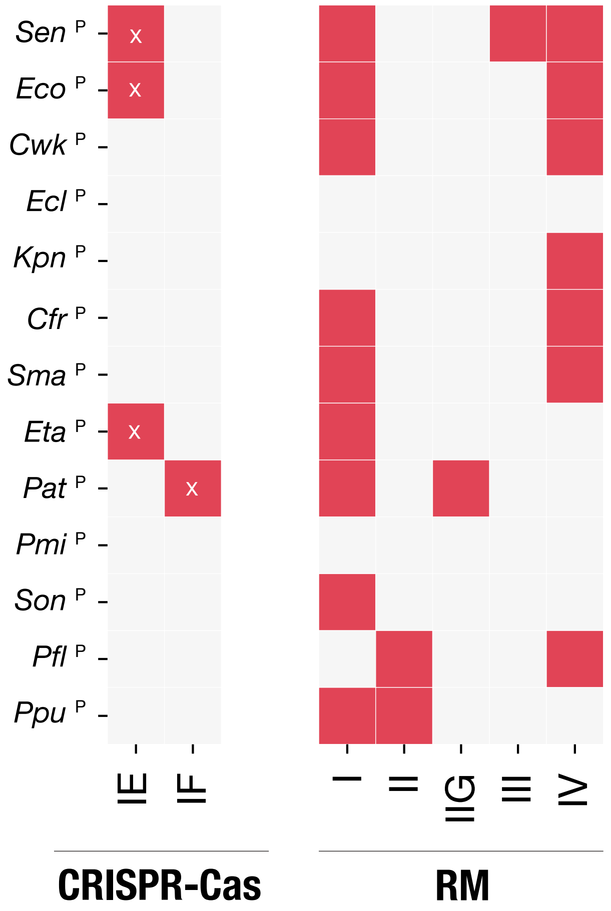


**Fig S4 | a,** Heatmap indicating presence or absence of CRISPR-Cas and restriction modification (RM) systems in the genomes of bacterial hosts used in this study, as detected by CRISPRCasTyper and DefenseFinder (Methods). Red indicates presence, and gray indicates absence of each specific subtype. Where hosts were found to contain CRISPR-Cas systems, the sequence of each spacer was queried against the sequence of all the phages in our collection to detect putative targeting events. A cross indicates that no spacers in the CRISPR arrays matched any of the plasmid-dependent tectiviruses in our collection. (Table S3)

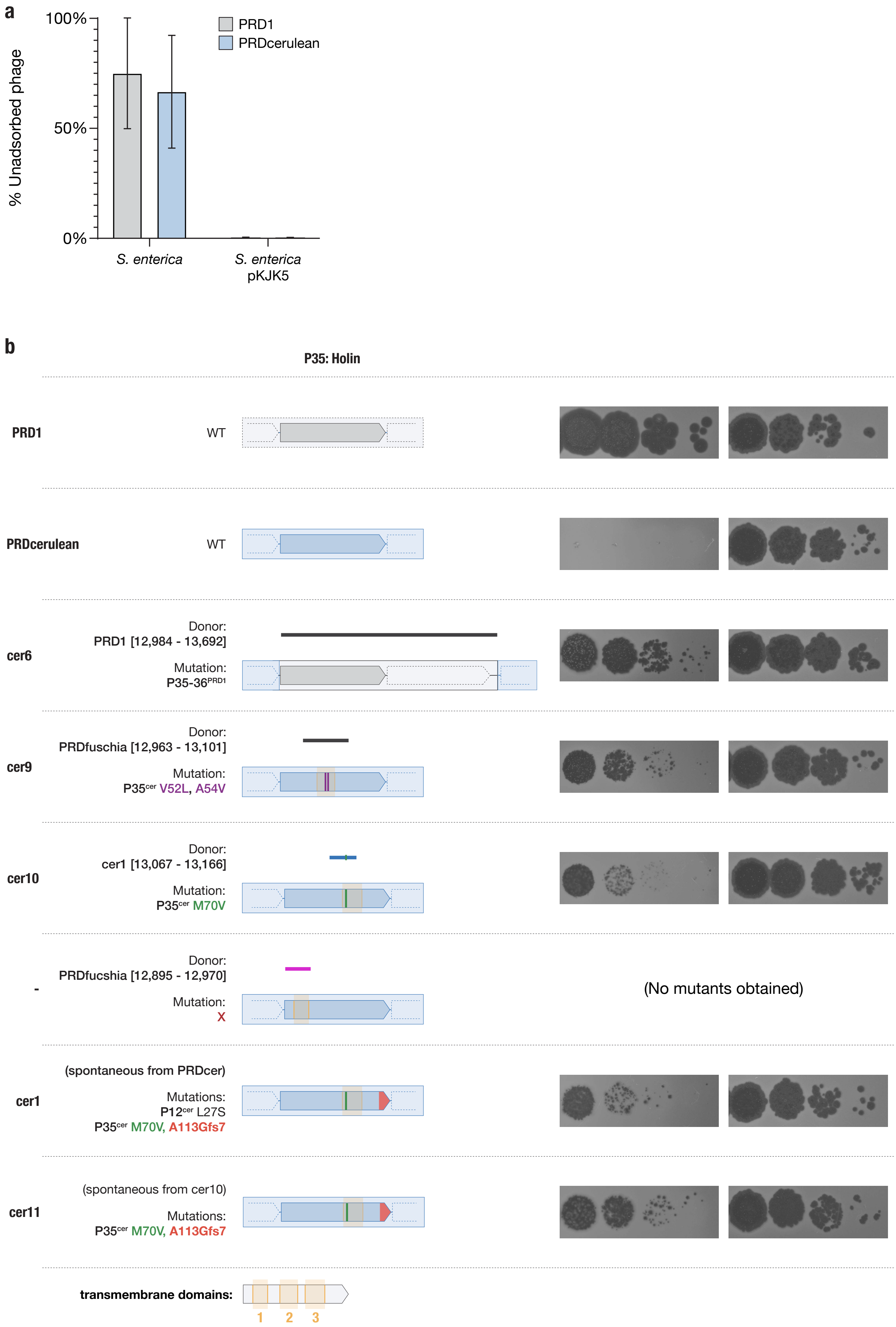
a
% Unadsorbed phage
100%
50%
0%
PRD1
PRDcerulean
S. enterica
S. enterica pKJK5
b
P35: Holin
PRD1
WT
PRDcerulean
WT
cer6
Donor:
PRD1 [12,984 - 13,692]
Mutation:
P35-36PRD1
cer9
Donor:
PRDfuschia [12,963 - 13,101]
Mutation:
P35cer V52L, A54V
cer10
Donor:
cer1 [13,067 - 13,166]
Mutation:
P35cer M70V
-
Donor:
PRDfucshia [12,895 - 12,970]
Mutation:
X
(No mutants obtained)
cer1
(spontaneous from PRDcer)
Mutations:
P12cer L27S
P35cer M70V, A113Gfs7
cer11
(spontaneous from cer10)
Mutations:
P35cer M70V, A113Gfs7
transmembrane domains:
1
2
3

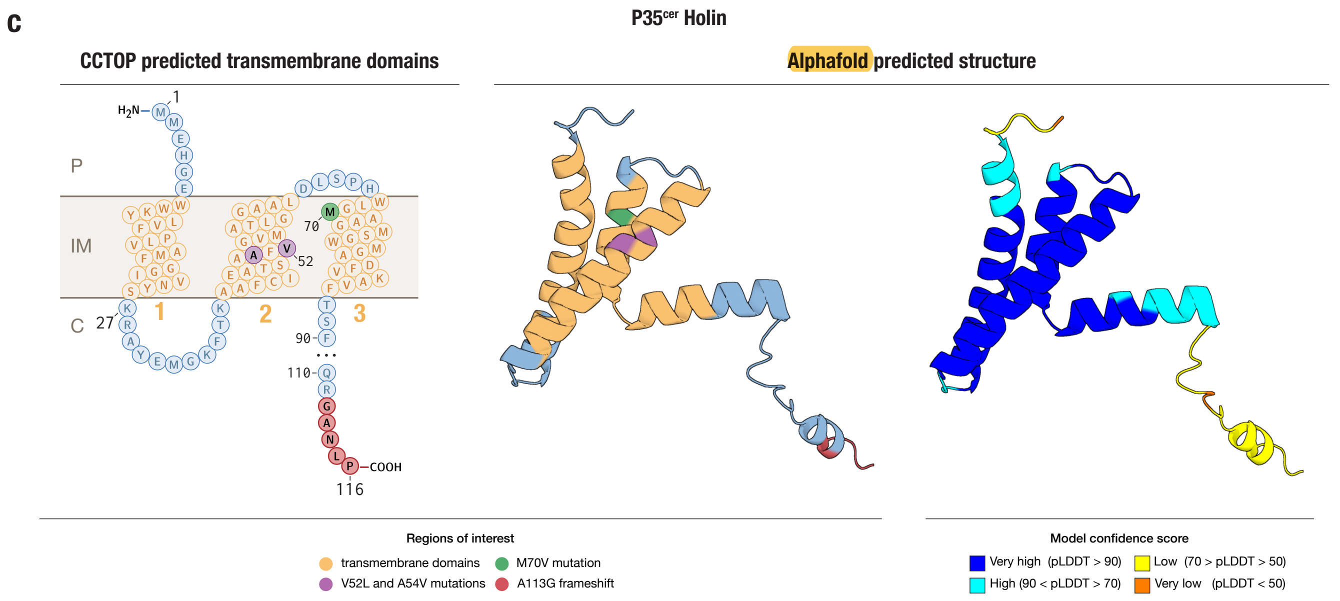
c
P35cer Holin
CCTOP predicted transmembrane domains
Alphafold predicted structure
P
IM
C
H2N
COOH
1
27
52
70
90
110
116
1
2
3
Regions of interest
transmembrane domains
M70V mutation
V52L and A54V mutations
A113G frameshift
Model confidence score
Very high (pLDDT > 90)
High (90 < pLDDT > 70)
Low (70 > pLDDT > 50)
Very low (pLDDT < 50)

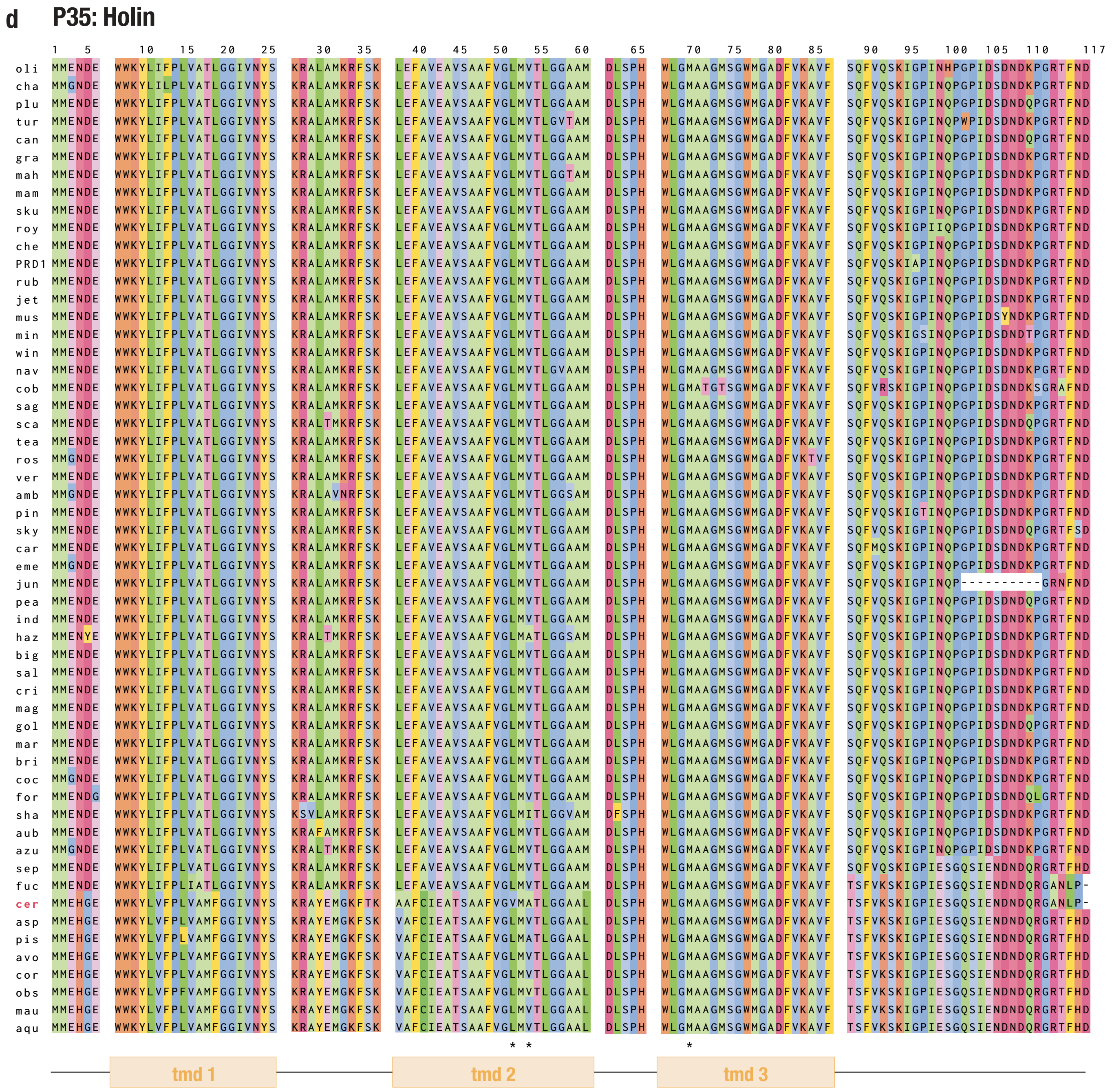
d
P35: Holin
1 5 10 15 20 25 30 35 40 45 50 55 60 65 70 75 80 85 90 95 100 105 110 117
oli
cha
plu
tur
can
gra
mah
mam
sku
roy
che
PRD1
rub
jet
mus
min
win
nav
cob
sag
sca
tea
ros
ver
amb
pin
sky
car
eme
jun
pea
ind
haz
big
sal
cri
mag
gol
mar
bri
coc
for
sha
aub
azu
sep
fuc
cer
asp
pis
avo
cor
obs
mau
aqu
tmd 1
tmd 2
tmd 3

**e** **P35: Holin mutants**

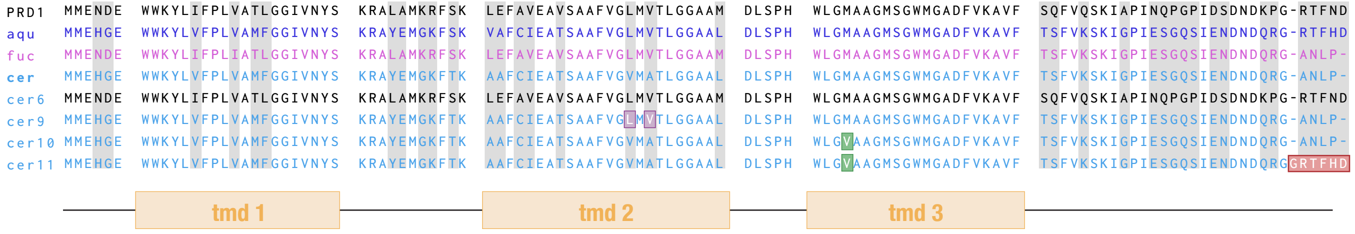


**Fig S5 | a,** Adsorption efficiency of PRD1 and PRDcerulean on *S. enterica* and *S. enterica* - pKJK5 (S. enterica[P]). Percentage of unadsorbed phage on *S. enterica* with no plasmid is shown as a negative adsorption control. There is no difference in adsorption between PRD1 and PRDcerulean (n=3) on the plasmid-bearing host. **b,** Detail of holin (P35) mutants and their effect in plaque formation. (Iconography as described in Fig. 5) Donor sequence used as recombination substrate for each of the phages is shown as a line above each mutant gene depiction, and the origin and precise coordinates of all the recombination substrates is given to the left. Recombination with TMD1 did not result in recombinant phages able to plaque on *S. enterica*[P] but recombinants of TMD2 & TMD3 did. **c,** To the left, a diagram of the predicted membrane topology of the holin protein (P35[cer]). It is predicted to have a short N-terminal periplasmic segment, 3 transmembrane domains (TMDs), and a longer disordered C-terminal region that extends into the cytoplasm. P = periplasm, IM = inner membrane and C = cytoplasm. Residues mutated in (b) are colored as indicated in the key. To the right, the AlphaFold predicted structure of P35[cer]. The structure is colored firstly by relevant region as previous, and secondly by model confidence score (pLDDT, a confidence score based on the local distance difference test), showing very high and high confidence in the membrane embedded section of the structure. This conformation suggests that the TMD2 and TMD3 mutations may be spatially proximal in the native protein structure. **d,** Holin protein multiple sequence alignment. Each row corresponds to a different isolate of our alphatectivirus collection (using abbreviated names). Vertical spaces have been added to highlight the location of the transmembrane domains, annotated at the bottom of the alignment.Asterisks mark the locations of the mutations of interest. **e,** Amino acid sequence of the holin mutants, columns in grey highlight non-conserved amino acid positions. PRDfuschia (fuc) and PRDcerulean (cer) share a distinct C-terminal end motif, but the differ in TMD1&2. Replacement of the full holin sequence (from PRD1 into PRDcerulean) and specific mutations in TMD2 and TMD3 in cer9 and cer10. respectively, recover the lysis phenotype. Finally, cer11, a large plaque mutant of cer10, reverts the C-terminal motif to the longer version, similar to that one of PRDaquamarine (aqu) and others, as shown in (d).

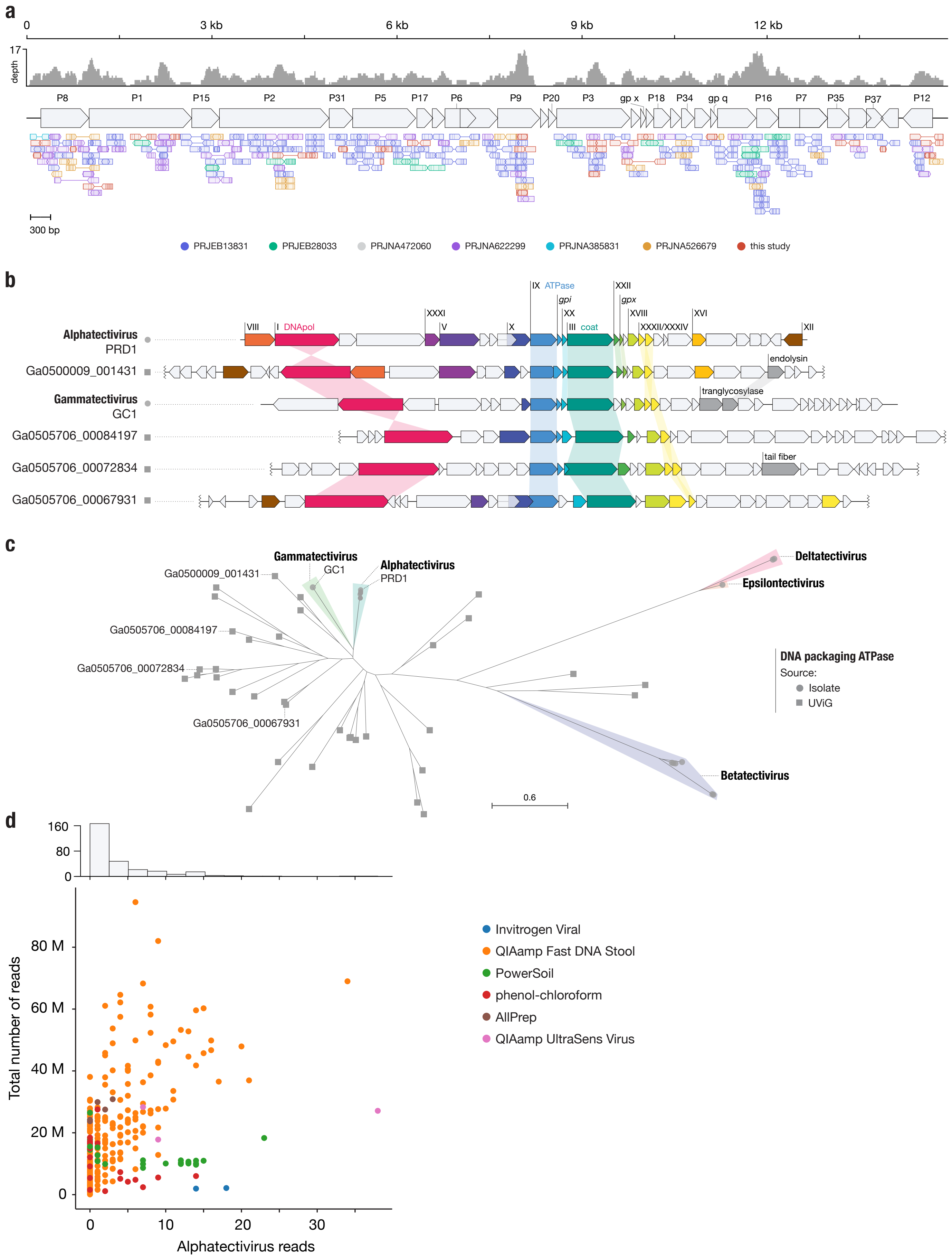


**Fig S6 | a,** Metagenomic reads mapped to regions of the PRD1 reference genome. ORFs are indicated with large gray arrows, and individual reads are represented by the smaller arrows and colored according to the dataset of origin (Fig. 6b), as described in the key. Direction of the arrows representing reads indicates the read direction, and lines between arrows indicate paired reads, with mismatches marked as vertical lines. **b,** Gene maps comparing Alphatectivirus PRD1 and Gammatectivirus GC1 against representative tectiviruses recovered from uncultivated viral genomes (UViGs). Colored genes represent homologs as detected by our protein models (HMMs) and shaded connectors represent proteins with >0.3 amino acid sequence identity. **c,** Maximum likelihood tree of the DNA packaging ATPase (terminase), including uncultivated tectiviruses (squares) and representatives of each genera of isolated tectiviruses (circles) **c,** Histogram and scatter plot of reads classified as being of alphatectiviral origin, against total number of reads in each metagenomic sample analyzed. Colors indicate different nucleic acid extraction methods as specified in the SRA metadata, full metadata of the datasets can be found in Table S2.
